# Multiomic analysis reveals that polyamines alter *G. vaginalis*-induced cervicovaginal epithelial cell dysfunction

**DOI:** 10.1101/2025.11.20.689523

**Authors:** Lauren Anton, Olha Kholod, Rutuja Phatate, Briana Ferguson, Kristin Klohonatz, Brittany A. Goods, Kristin D. Gerson

**Affiliations:** Center for Women’s Health and Reproductive Medicine, Department of Obstetrics and Gynecology, Perelman School of Medicine at the University of Pennsylvania, Philadelphia, PA 19104, USA; Thayer School of Engineering at Dartmouth College, Hanover, NH, USA; Department of Molecular and Systems Biology, and Program in Quantitative Biomedical Sciences at Dartmouth College, Hanover, NH, USA; Department of Microbiology, Perelman School of Medicine at the University of Pennsylvania, Philadelphia, PA, 19104

## Abstract

An anaerobe-dominant, *Lactobacillus*-deplete cervicovaginal microbiome is associated with adverse reproductive outcomes. *Gardnerella vaginalis*, a cervicovaginal anaerobe, alters cervicovaginal epithelial cell function, resulting in immune activation and barrier breakdown. Host-microbial mechanisms inducing this epithelial dysfunction remain unknown. We show microbe-specific alterations in cervicovaginal epithelial cell metabolite profiles where *G. vaginalis*, but not *Lactobacillus crispatus*, increases polyamine biosynthesis. Pretreatment with polyamines (putrescine, spermidine and spermine) globally shifts *G. vaginalis*-induced transcriptomic profiles. Alterations in enzyme transcripts responsible for polyamine synthesis and catabolism provide evidence that *G. vaginalis* modifies polyamine biosynthesis. Polyamine-mediated transcriptomic changes include genes related to bacterial defense, inflammation, and epigenetic processes. Polyamines mitigate *G. vaginalis*-induced inflammatory responses through reduction of cytokines/chemokines and matrix metalloproteinases. *In vitro* transcriptional signatures positively correlated to existing human datasets. The ability of cervicovaginal metabolites to alter microbe-mediated changes in epithelial cell function suggests that metabolite-microbe interactions are critical mediators of epithelial defense against a *Lactobacillus*-deplete microbiota.

## Introduction

The female lower reproductive tract contains a complex ecosystem of microorganisms, metabolites, immunomodulatory components and host cells, which are essential to maintaining homeostasis and reproductive health. Numerous studies have characterized the vaginal microbiome in non-pregnant and pregnant individuals and have found associations with health and disease.^1–4^ Though less complex than microbial ecosystems at other body sites such as the gut, mouth, or skin, bacterial diversity and instability of the vaginal microbiome have been associated with several disease pathologies, including infertility and adverse gynecological and obstetrical health outcomes.^5^ The vaginal microbiome has been broadly divided into *Lactobacillus-*dominant (low diversity) and *Lactobacillus*-deplete (high diversity) classifications. More granular classification schemes separate communities into state types (CSTs) based on predominant bacteria.^1,2,6^ Clinical studies have shown that a *Lactobacillus*-dominant vaginal microbiome, most often classified by high levels of *Lactobacillus crispatus* (*L. crispatus*), is associated with overall reproductive health and is protective against vaginal infection.^7,8^ This *L. crispatus*-mediated protection is largely due to the high production of lactic acid,^9,10^ immunomodulatory anti-microbial peptides,^11,12^ bacteriocins,^13^ and hydrogen peroxide,^14^ all of which create an unfavorable environment for vaginal pathogens.

Vaginal microbial communities with a high abundance of anaerobes and a low abundance of *lactobacilli* are associated with a high risk for bacterial vaginosis (BV), ^15,16^ sexually transmitted infections (i.e., HIV and HPV),^17–19^ and adverse pregnancy outcomes (i.e., miscarriage and preterm birth).^4,20^ This microbial dysbiosis is often characterized by an overgrowth of strict and facultative anaerobes, including *Prevotella*, *Atopobium*, and *Mobiluncus* subspecies, and most predominantly *Gardnerella vaginalis* (*G. vaginalis)*. These vaginal bacteria alter the microenvironment, leading to an increase in pH, inflammatory mediators,^21^ mucin degrading enzymes (e.g., sialidases),^22^ and select metabolites (e.g., short chain fatty acids and biogenic amines).^23–25^ Despite the associated risk, *G. vaginalis* has been detected in more than 50% of women who do not show clinical symptoms of BV.^15,26,27^ While studies using advanced molecular technologies including 16s rRNA sequencing, metagenomics, metatranscriptomics, and proteomics have helped to further refine the classification of *Gardnerella* subspecies (clades) and their associated virulence factors,^28^ elucidating the specific biological mechanisms through which these bacteria exert their functional effects are essential to the future development of targeted therapeutic strategies aimed at mitigating adverse gynecological and reproductive outcomes.

Vaginal bacteria interact with cervicovaginal epithelial cells and/or their cell surface receptors to regulate host biological functions, including immune responses, metabolism, cellular signaling, and epithelial barrier function. These host-microbe interactions are critical to the defense and maintenance of the cervicovaginal epithelial barrier. Previous studies by our group and others have shown that *G. vaginalis* decreases epithelial barrier integrity and activates a TLR2/ NF-kB-mediated innate immune response in cervical and vaginal epithelial cells.^29–32^ Additionally, RNA sequencing of cervical cells exposed to *G. vaginalis* showed an upregulation of innate immune genes, including NF-kB-mediated cytokines and anti-microbial peptides.^33^ However, the molecular pathways mediating this host-microbial immune activation are complex and remain undefined.

Metabolites can act as inert byproducts of cellular metabolism or play key roles in modifying diverse physiological processes. Recent studies, most predominantly in the gut, have identified that metabolites produced by both bacteria and host cells play an important role in regulating epithelial cell function.^34–37^ Previous studies have also shown that select vaginal metabolites correlate with microbial dysbiosis in pregnancy, and that metabolomic models can be used to identify individuals at greatest risk of preterm birth among those with similar microbiome-related risk.^25,38,39^ Overgrowth of *G. vaginalis* is associated with increased abundance of biogenic amines, including putrescine, cadaverine, and tyramine, which contribute to the clinical features of increased vaginal pH and malodor seen in BV.^40^ These biogenic amines reduce *L. crispatus* growth and decrease lactic acid production, further driving vaginal dysbiosis.^41^ Polyamines are a class of biogenic amines that includes putrescine and its downstream metabolites spermidine and spermine. In the gut, polyamines have been shown to alter epithelial cell function affecting proliferation, migration, apoptosis, and cell-cell tight junctions which are important for epithelial barrier strength and homeostasis ^34,42–44^. Importantly, polyamines can alter the immune response or mitigate inflammatory damage as part of the immunometabolome.^45^ Spermidine induces an anti-inflammatory macrophage phenotype^46,47^ and regulates differentiation of T, B and natural killer cells, ^48–51^ while spermine mitigates proinflammatory cytokines in monocytes.^52^

Despite evidence that polyamines are altered by the vaginal microbiome and can impact epithelial and immune cell function, the role of polyamines in regulating microbe-mediated immune responses in the cervicovaginal space is unknown. To address this key knowledge gap, we leveraged a multi-omics discovery-based approach to investigate the pathways activated in cervicovaginal epithelial cells exposed to *G. vaginalis,* and to determine how vaginal polyamines alter this response. Specifically, cervicovaginal epithelial cells exposed to *L. crispatus* or *G. vaginalis* resulted in microbe-specific alterations in metabolite profiles and pathways including polyamine biosynthesis. Using RNA-sequencing, transcriptional changes and activated functional pathways were identified in *G. vaginalis*-exposed cervicovaginal cells and, secondarily, we found that these changes were altered by pretreatment with polyamines. Through WCGNA analysis, we identified the top genes associated with polyamine pretreatment in *G. vaginalis*-exposed cells, many of which are known regulators of epithelial cell function. Focusing on the cervicovaginal host immune response, we used Luminex to profile polyamine alterations to *G. vaginalis*-induced cytokine production. Lastly, we compared our *in vitro* findings to human transcriptomic datasets from women with a Lactobacillus-deplete microbiome. Overall, our findings provide evidence of microbe-specific alterations to the cervicovaginal epithelial cell metabolome and show that *G. vaginalis*-induced changes to epithelial cell function can be mediated by polyamines.

## Materials and Methods

### Eukaryotic cell culture

Ectocervical (Ect/E6E7, ATCC# CRL-2614) (Ecto), endocervical (End1/E6E7, ATCC# CRL-2615) (Endo) and vaginal (VK2/E6E7, ATCC# CRL-2616) (VK2) human epithelial cell lines (American Type Culture Collection, Manassas, VA) were cultured in Keratinocyte-Serum Free Media (KSFM) supplemented with 0.1 ng/mL epidermal growth factor and 50 μg/mL bovine pituitary extract (Gibco, Life Technologies), 100 U/mL penicillin, and 100 μg/mL of streptomycin at 37°C in a 5% CO_2_ humidified incubator.

### Bacterial cell culture

Human clinical isolates of *L. crispatus* (LC, ATCC 33197) or *G. vaginalis* (GV, ATCC 14018), were obtained from the American Type Culture Collection (Manassas, VA). *G. vaginalis* was grown on Tryptic Soy Agar with 5% Sheep Blood plates (Hardy Diagnostics) and *L. crispatus* was grown on De Man, Rogosa and Sharpe agar (Fisher Scientific); both strains were grown in New York City III (NYCIII) broth. Bacteria were grown at 37°C in an anaerobic glove box (Coy Labs, Grass Lake, MI).

For each experiment the following bacterial growth protocol was followed: *L. crispatus* and *G. vaginalis* glycerol stocks were streaked on agar plates, as well as into broth tubes and grown overnight. The broth starter cultures were diluted to an optical density of 0.2 and then used to inoculate 20ml working cultures, which were grown for 20 hours (*G. vaginalis*) to 48 hours (*L. crispatus*) prior to use in experiments. Bacterial densities of the working cultures were estimated the day of the experiment based on optical density readings at 600 nm using an Epoch2 plate reader (Biotek, Winooski, VT), and the appropriate volume was centrifuged at 13,000 x *g* for 3 min. The bacterial pellets were resuspended in the appropriate cell culture media without antibiotics and added to epithelial cells at 10^7^ CFUs/well. Precise bacterial densities of the working cultures were determined by plating serial dilutions of the working cultures and counting CFUs. For all experiments, reported bacterial densities are +/- 0.5 log of the noted bacterial density (CFU/well).

### Bacterial and polyamine exposure experiments

Ecto, Endo and VK2 epithelial cells were plated at 1.5 x 10^5^ cells/well in twenty-four well plates containing KSFM without antibiotics. The next day, cells were pretreated for 4 hours with or without polyamines, putrescine (PUT, 4mM), spermidine (SPD, 400uM) or spermine (SPM, 400uM) prior to exposure to *G. vaginalis* or *L. crispatus* (10^7^ CFU/well) for an additional 24 hrs. Final polyamine concentrations were chosen based on preliminary dose response experiments in Endo cells (**Figure S1**). Cell culture media was collected for cell death assays (**Figure S2**), metabolomics analysis and Luminex and/or the cells were collected in Trizol (Invitrogen, Thermo-Fisher Scientific) for RNA extraction and RNA sequencing.

### Metabolomics data generation

Cell culture media collected from the experiments described above were sent to Metabolon, Inc. (Morrisville, NC) for global metabolomics analysis. To allow for capture of all small molecules including diverse metabolites, proteins were removed via precipitation with methanol. The resulting extracts were divided into multiple fractions: two for analysis by two separate reverse phase (RP)/UPLC-MS/MS methods with positive ion mode electrospray ionization (ESI), one for analysis by RP/UPLC-MS/MS with negative ion mode ESI, and one for analysis by HILIC/UPLC-MS/MS with negative ion mode ESI. The details of this global metabolomics platform have been described previously.^53,54^ This platform allows for the identification of a broad range of compounds with different physiochemical properties such as mass, charge, chromatographic separation, and ionization behavior. Controls included pooled sample matrix, extracted water samples (process blanks), and a cocktail of QC standards spiked into analyzed samples to protect against process variability/contamination. Raw data was extracted, peak-identified and QC processed using proprietary Metabolon developed software. Peaks were then quantified using area-under-the-curve. Compounds were identified by comparison to a Metabolon curated library of purified standards containing retention time/index (RI), mass to charge ratio (*m/z)*, and fragmentation data.^55,56^ Biochemical identifications were based on three criteria: retention index, accurate mass match to the library (+/- 10 ppm), and the MS/MS forward and reverse scores (ions) between experimental data and authentic standards. Metabolomics data are provided in **Table S1**.

### Metabolomics data analysis

The data from Metabolon were imported into the MetaboAnalyst platform for further analysis.^57^ Analyses for each cell type (Ecto, Endo, and VK2) were performed separately. The data were normalized by log-transformed with base 10. We used the Human Metabolome Database (HMBD) for pathway annotations.^58^ The enrichment ratio was calculated as the ratio of observed hits to expected hits. To select the top 15 pathways of interest, we transformed outputted FDR values as -log10P, sorted the list of pathways by -log10P in decreasing order, and then selected the top pathways that have at least five representative hits in our dataset.

To generate correlation plots for putrescine, spermidine, and N(1)-acetylspermidine, we uploaded data from Metabolon into the statistical analysis (one factor) tab in the MetaboAnalyst platform. We created separate analyses for each cell type (Ecto, Endo, and VK2), including all conditions (Non-treated (NT), *L. crispatus*, and *G. vaginalis*). We applied the default filtering parameters, which exclude 10% of variables based on interquartile range (IQR). The data were then normalized by log-transformed with base 10. Under univariate analysis, we selected the “Pattern search” option and specified putrescine, spermidine, and N(1)-acetylspermidine as features of interest. We performed Pearson correlation analysis to identify metabolites that correlated with these compounds. The top 25 correlated compounds were visualized as a bar plot, with the x-axis representing the correlation coefficient and the y-axis representing metabolites that positively or negatively correlated with the metabolite of interest (**Figure S3**).

### Transcriptomics data generation

After exposure to *G. vaginalis* +/- PUT, SPD, or SPM, Ecto, Endo and VK2 cells were prepared for RNA-sequencing. All samples were processed and sequenced at the Children’s Hospital of Philadelphia (CHOP) High Throughput Sequencing Core. RNA-sequencing libraries were generated according to manufacturer’s instructions for the CORALL Total RNA-Seq Library Prep kit (Lexogen GmbH). Unique Dual Indexes Primer Pair Set were incorporated for multiplexed high-throughput sequencing. The resulting RNA was assessed for size distribution, concentration and RNA integrity (RIN) using the Bioanalyzer High Sensitivity DNA Kit (Agilent Technologies) with all samples having RIN values >9. Input RNA (400 ng) was depleted of ribosomal RNA with RiboCop for Human/Mouse/Rat plus Globin. Library generation, performed according to the manufacture’s protocol (Lexogen NGS Services, Vienna, Austria), was initiated by random hybridization of Displacement Stop Primers to the RNA template. These primers contain partial Illumina-compatible P7 sequences. Reverse transcription extends each DSP to the next DSP where transcription is effectively stopped. This stop prevents spurious second strand synthesis and thus maintains excellent strand specificity and seamlessly integrates Unique Molecular Identifiers (UMIs). UDIs indexes were added during the PCR amplification step, in which complete adapter sequences required for cluster generation on Illumina instruments were also added. All nucleic acid purification steps used AMPure XP magnetic beads (Beckman Coulter).

The resulting libraries were pooled, diluted to 2 nM using 10 mM Tris-HCl, pH 8.5, denatured, and loaded onto an S2-200 (PE100) flow cell on an Illumina NovaSeq 6000 (Illumina, Inc.) according to the manufacturer’s instructions. De-multiplexed and adapter-trimmed sequencing reads were generated using Illumina bcl2fastq.

Transcript quantification from RNA-seq data was performed using Salmon and release 44 (GRCh38.p13) of the human genome. QC analyses were done with Fastqc for the raw fastq files. Several Bioconductor (v3.17) packages in R (v4.32) were used for subsequent steps. The transcriptome count data was annotated and summarized to the gene level with tximeta and further annotated with biomaRt. Normalizations and statistical analyses were done with DESeq2. The raw RNA-seq data was submitted to Gene Expression Omnibus (accession #GSE311352).

### Differential expression analysis and pathway analysis

We performed differential expression (DE) analysis using the DESeq2 R package (v1.12.3) to identify genes with significant expression changes in response to polyamine+*G. vaginalis* treatment. Pairwise comparisons were made between polyamine+*G. vaginalis* and non-treated+*G. vaginalis* conditions for each cell type (Ecto, Endo, and VK2). Genes with a log_2_ fold change (logFC) ≥ 2 and an adjusted p-value < 0.05 were considered differentially expressed.

Functional annotation of upregulated genes was performed using Metascape.^59^ We selected top pathways annotated as Gene Ontology Biological Processes (GO_BP) and transformed the outputted false discovery rate (FDR) values as -log10P. To avoid including categories supported by only a few genes, we examined the hierarchical clustering of pathways in the Metascape output and retained the top pathways from each summary category that were represented by at least five genes in our dataset. For example, if six of our upregulated genes mapped to a GO_BP term (e.g., 6 of 37 annotated genes), this pathway was included in the bar plot; however, if only two genes mapped to a term (e.g., 2 of 43), it was excluded.

### Weighted gene co-correlation (WGCNA) analysis

To obtain modules of genes with similar expression profiles across our treatment and bacterial exposure phenotypes, we performed WCGNA analysis.^60^ The input genes for WCGNA were selected by obtaining the union of differentially expressed genes derived from DE analysis and by setting a threshold of 0.5 on log_2_ fold change and 0.00001 on the p.adj values. The soft threshold parameter was chosen to be 9 (corresponding to approximate scale-free topology) and a signed network of the genes was generated on DESeq2 normalized counts. The original 27 modules were merged into final 9 modules (including module grey). A module trait relationship heatmap was generated separately for each cell type and treatment_bacterial exposure status phenotypes. Functional enrichment analysis of the genes contained in this module was performed using the CluGO app on Cytoscape 3.10.3 version. Enrichment terms were pulled from GO, KEGG and Reactome.

### Merged network analysis

The WCGNA topological overlap matrix (TOM) was subset (subTOM) for genes contained in module darkgreen and a 90^th^ percentile threshold was implemented for selecting edges. The similarity values contained within subTOM were used as edge weights. Node attributes of intramodular connectivity and module membership (kME) measure were calculated based on normalized counts, adjacency and module eigengene values for genes contained in module darkgreen. To select the top genes correlated with treatment_*G. vaginalis* (GV) exposure, the metadata was modified by binarizing exposure_status into exposure (GV) and no exposure (noGV) and treatment_type was grouped into four levels: non-treated (NT), PUT, SPD and SPM. A model matrix design of exposure_status + treatment_type + treatment_type:exposure_status was implemented to obtain correlation values. Top 50 genes with positive kME and the highest correlation with exposure_statusGV:treatment_typePUT, exposure_statusGV:treatment_typeSPD and exposure_statusGV:treatment_typeSPM were selected and a separate network of these genes for each treatment was built using Cytoscape 3.10.3 version. Top 10 DE genes were selected based on highest average of log_2_ fold change values between the NT+GV and polyamine+GV DE lists from all three cell lines. These 10 genes were highlighted in yellow on the network and plotted on the heatmap in figure 3.

**Figure 1.**
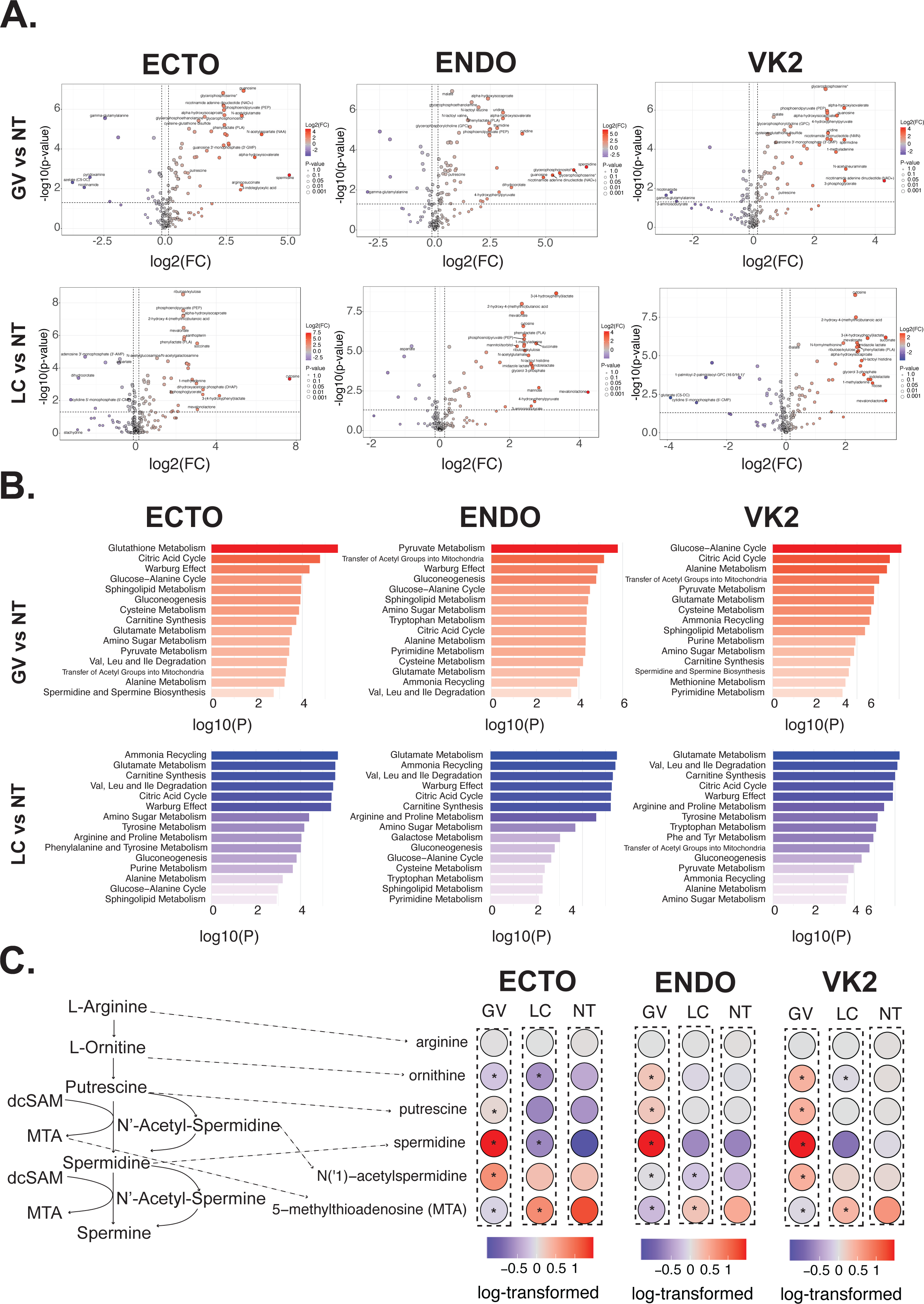
Metabolite profiles are altered across Ectocervical (Ecto), Endocervical (Endo), and Vaginal (VK2) cells in response to *L. crispatus* (LC) or *G. vaginalis* (GV). (A) Volcano plots displaying differentially abundant metabolites between GV vs NT (top) and LC vs NT (bottom). Red and blue dots represent increased and decreased metabolites, respectively. Top 20 metabolites are labeled. (B) Bar plots showing pathways enriched in GV (red) and LC (blue) exposures relative to non-treated samples (FDR = 0.05). The x-axis represents -log10(p) values. (C) Schematic representation of polyamine metabolites (left) and corresponding dot plots in Ecto, Endo, and VK2 cell types (right). Raw values were log-transformed. Red colored dots indicate highest metabolite abundance. Significant metabolites marked with asterisks. *p<0.05

**Figure 2.**
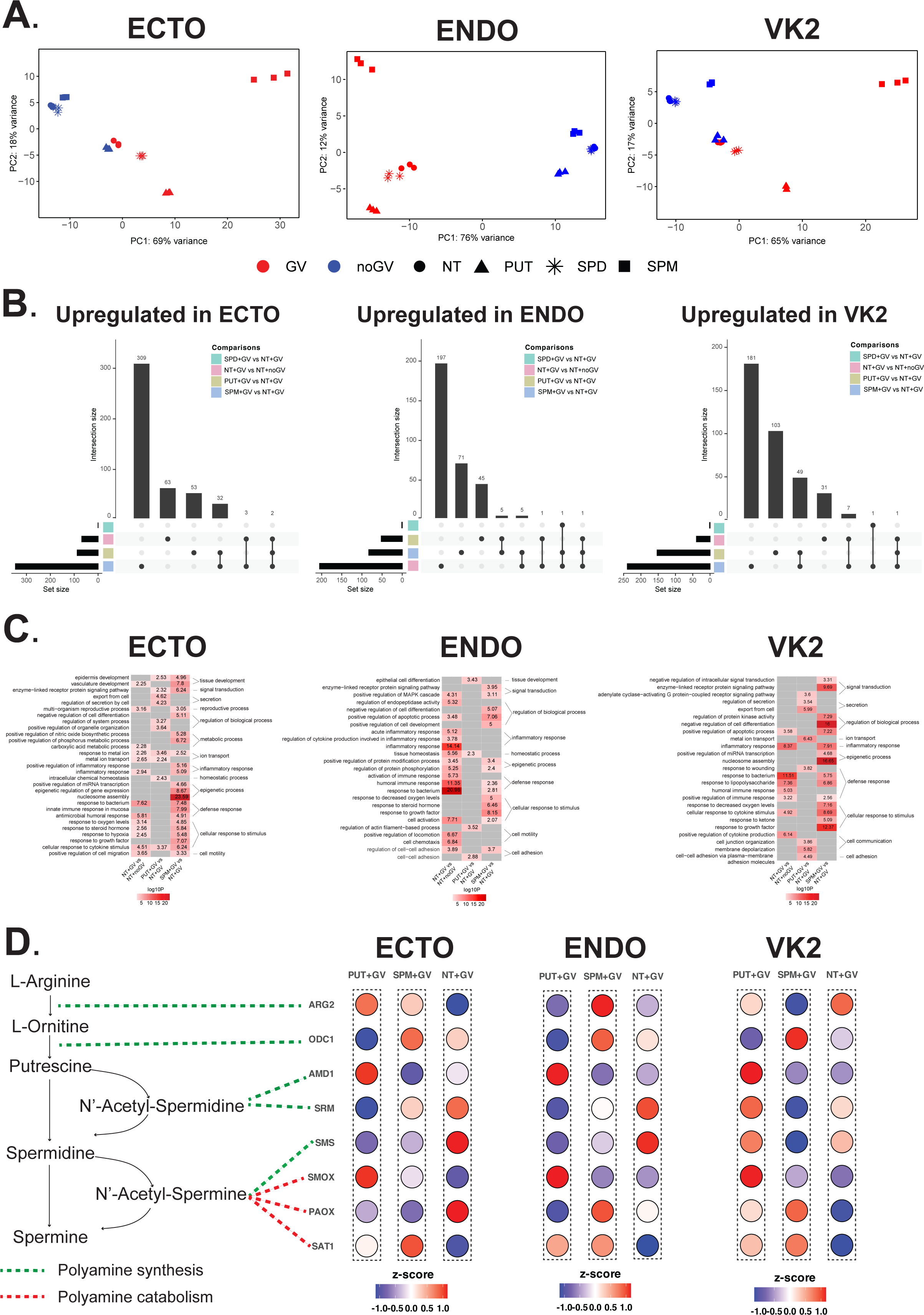
Transcriptomic analysis of Ectocervical (Ecto), Endocervical (Endo) and Vaginal (VK2) cells reveals global transcriptomic shifts that are impacted by exposure to *G. vaginalis* (GV) in the presence of putrescine (PUT), spermidine (SPD) or spermine (SPM). (A) PCA plots along the first two principal components shown for Ecto, Endo, and VK2 cells with or without polyamine pretreatment (NT, PUT, SPD, or SPM) and colored by GV (red) or no GV (blue) exposure. (B) Upset plots show overlapping genes for each indicated comparison by upregulated gene lists, where input gene lists were filtered based on significance (padj < 0.05) and fold change (FC > 2). (C) Heatmaps show top pathways derived from a unique set of upregulated genes in NT+GV vs NT+noGV, PUT+GV vs NT+GV and SPM+GV vs NT+GV comparisons from each cell type. Gene sets were extracted from upset plots in Figure 2B. (D) Dot plots in Ecto, Endo, and VK2 cells showing genes encoding enzymes that regulate polyamine biosynthesis. Results are shown as log_2_FC values between conditions of interest. Red dots indicate upregulated genes and blue dots represent down-regulated genes.

**Figure 3.**
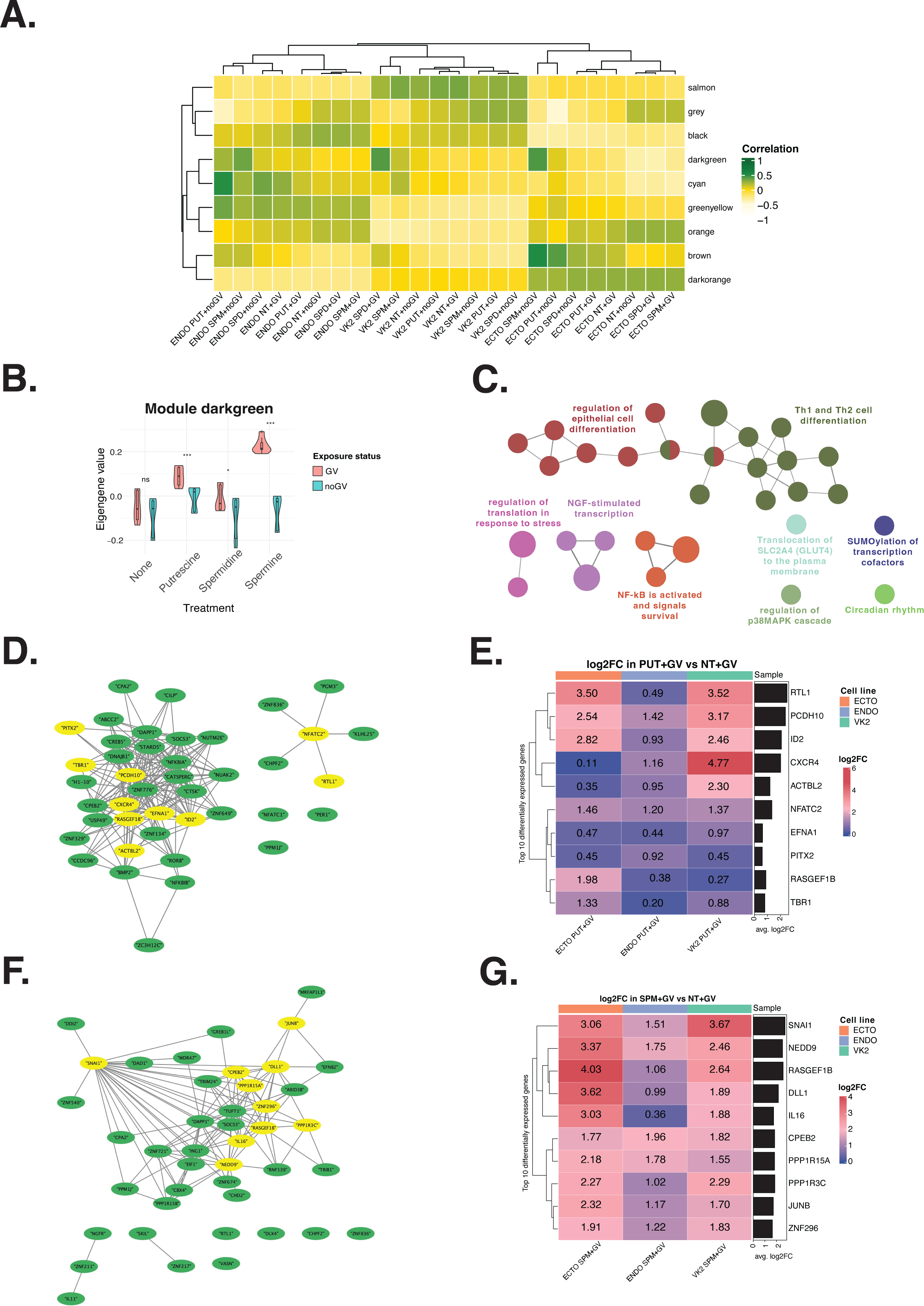
WCGNA analysis identifies modules correlated with putrescine (PUT) and spermine (SPM) pretreatment in *G. vaginalis* (GV)-exposed Ectocervical (Ecto), Endocervical (Endo) and Vaginal (VK2) cells. (A) Module-trait relationship heatmap showing the correlation between each sample group and the module eigenvector. P-values associated with the correlation coefficient are represented in parentheses. (B) Polyamine pretreatment and bacterial exposure conditions from all cell types were aggregated to identify modules with consensus signatures. Module darkgreen eigengene value for all polyamine and GV exposures were significantly lower compared to polyamines with no GV exposure. We picked this module for further investigation. (C) Gene enrichment results for genes from module darkgreen. (D) Network of top 50 genes associated with PUT+GV exposure across cell types. Top 10 genes based on average fold change between NT+GV and PUT+GV are highlighted in yellow. (E) Heatmap showing the fold change value between NT+GV and PUT+GV across all cell types for the 10 genes highlighted in yellow in Figure 3D. (F) Network of top 50 genes associated with SPM+GV exposure across all cell types with top 10 genes highlighted in yellow. Top 10 genes based on average fold change between NT+GV and SPM+GV are highlighted in yellow. (G) Heatmap showing the fold change value between NT+GV and SPM+GV across all cell types for the 10 genes highlighted in yellow in Figure 3F.

### Luminex data generation

Ecto, Endo and VK2 cells were cultured and treated with live bacteria +/- polyamines as stated above. A 41-plex cytokine/chemokine (HCYTMAG-60K-PX41) and an MMP 5-plex (HMMP2MAG-55K) human magnetic bead Luminex panel (EMD Millipore, Billerica, MA) was run by the Human Immunology Core at the University of Pennsylvania on Ecto, Endo and VK2 cell culture media after 24 hours of treatment with *L. crispatus* (n=3) or *G. vaginalis* (n=3) live bacteria. All samples were run in duplicate, per the manufacturer’s protocol on the FLEXMAP3D Luminex platform (Luminex, Austin, TX). Absolute quantification in pg/mL was obtained using a standard curve generated by a five-parameter logistic (5PL) curve fit using Xponent 4.2 software (Luminex).

### Luminex data analysis

Luminex multiplex immunoassay data were processed and analyzed in R (v4.5.0). PCA was performed using the R packages dplyr (v1.0.10), Biobase (v2.32.0), PCAtools (v2.5.13), and ggplot2 (v3.4.3). Input data consisted of count tables of observed concentrations for 46 analytes, together with sample metadata including cell type, polyamine treatment, and *G. vaginalis* exposure. Each biological replicate was measured in triplicate. Analytes with zero values across all biological and technical replicates within a given cell type were removed, yielding 36 analytes for Ecto, 35 for Endo, and 37 for VK2. PCA was conducted with a variance filter threshold of 0.5 to exclude low-variance features. The variance explained by each principal component was extracted from the PCA model. PCA scores were merged with sample metadata for visualization in the PC1–PC2 space. Data points were colored by *G. vaginalis* (GV) exposure (red = present, blue = absent) and shaped by polyamine treatment (circle = Non-Treated (NT), triangle = PUT, square = SPM, cross = SPD). Axes were annotated with the proportion of explained variance, and a fixed coordinate ratio was applied to preserve relative distances.

### Two-way analysis of variance (ANOVA) for Luminex data

Statistical analyses were performed in R using stats (v3.6.2) and dplyr (v1.0.10). Median fluorescence intensity (MFI) values were used instead of observed concentrations. To stabilize variance and approximate normality, MFIs were log10-transformed prior to analysis. The dataset comprised 46 analytes for each combination of cell type, polyamine treatment, and GV exposure, with three technical replicates per biological replicate. Categorical variables (cell type, treatment and GV exposure) were converted to factors. For each analyte within each cell type, a two-way ANOVA was fit with the model: *MFI* ∼ *Treatment* * GV. This model tested main effects of Treatment and GV, as well as their interaction. Pairwise differences between treatment–GV combinations were evaluated using Tukey’s Honest Significant Difference (TukeyHSD) test applied to the interaction term (Treatment: GV). TukeyHSD results included estimated mean differences, 95% confidence intervals, and adjusted p-values, correcting for multiple comparisons. Analyses were restricted to analyte–cell type combinations with ≥2 data points. For each valid combination, results were compiled into a summary table containing CellLine, Analyte, Comparison, estimated difference (Diff), confidence intervals (LowerCI, UpperCI), and adjusted p-value (P_value) **(Table S2**). Comparisons of primary interest are shown in **Table S3**: NT+noGV vs. NT+GV, PUT+GV vs. NT+GV, SPD+GV vs. NT+GV, and SPM+GV vs. NT+GV.

Analytes not significant in any of these comparisons were excluded.

### Comparison of transcriptional signatures between *in vitro* and *in vivo* studies

To assess whether transcriptional responses in our Ecto, Endo, and VK2 cells recapitulate gene signatures previously associated with either a *Lactobacillus*-dominant microbiome or preterm birth, we quantified the overlap between our *in vitro* experimentally derived genes and published *in vivo* reference sets using Fisher’s exact test. Reference gene lists were curated from transcriptomic studies, including Berard et al. ^24^ and Wikström et al.^61^ Experimental gene sets consisted of genes upregulated in each of the tested conditions (NT+GV, NT+noGV, PUT+GV, PUT+noGV, SPM+GV, and SPM+noGV).

For every pairwise comparison between experimental and reference datasets, we computed a 2×2 contingency table based on the number of shared genes, genes unique to each list, and a fixed background of 20,000 protein-coding genes. Fisher’s exact test was applied to each table, and the resulting p-values were transformed to –log_10_ values to facilitate comparison across conditions.

To aid interpretation, p-values were additionally converted to categorical significance labels, with one, two, or three asterisks denoting increasing levels of statistical significance. These labels were overlaid on the heatmap to highlight robust enrichments.

Heatmaps were generated using the ComplexHeatmap package in R. Row and column clustering were disabled to preserve the predefined ordering of experimental conditions and reference datasets. The resulting visualization provides a direct comparison of how strongly our *in vitro* experimental gene signature aligns with established microbiome or preterm birth associated signatures.

### Code and data accessibility

All processed data files are available as supplemental files with this manuscript. Raw transcriptomics data can be found on GEO, metabolomics data and Luminex data as a supplemental file. All code used to analyze data is available on the Goods Lab GitHub (https://github.com/Goods-Lab/Preterm-multiomics-analysis).

### Declaration of generative AI and AI-assisted technologies

During the preparation of this work, ChatGPT (versions 4o and 5) was used to aid in debugging R code. After using this tool or service, the author(s) reviewed and edited the content as needed and take(s) full responsibility for the content of the publication.

## Results

### Metabolite profiles are differentially modified by exposure to *L. crispatus* and *G. vaginalis*

Given evidence that the vaginal microbiome can directly modulate cell metabolism, we first sought to test the hypothesis that exposure of cervicovaginal cells to either *L. crispatus* or *G. vaginalis* leads to differential metabolite profiles. We exposed Ecto, Endo, and VK2 epithelial cells to live strains of *G. vaginalis* or *L. crispatus* for 24hrs, then collected supernatants for untargeted metabolomic analysis. After quality control and normalization, we performed differential metabolite analysis for Ecto, Endo, and VK2 cells. We found that cervicovaginal epithelial cell exposure to *G. vaginalis* or *L. crispatus* generated distinct metabolic signatures as compared to each other and relative to non-treated conditions (**Figure 1A**). While a select few metabolites were unique to each cell type, most differentially detected metabolites were consistently increased across all three cell types in response to either *G. vaginalis* or *L. crispatus* exposure (**Figure S3A**). Metabolites that were increased in response to *G. vaginalis* exposure across all cervicovaginal cell types included polyamines (putrescine and spermidine) as well as purine and pyrimidine metabolites (guanosine, uridine, cytidine, thymidine), and metabolites involved in energy metabolism through glycolysis and gluconeogenesis (**Table S4**). In contrast, metabolites related to succinate and lactate metabolism were highly produced in *L. crispatus*- exposed cells (**Figure 1A, Table S4**).

Next, we aimed to investigate the underlying metabolic pathways that were impacted by bacterial exposure. We conducted enrichment analysis on differentially detected metabolites using the Human Metabolome Database (HMDB) (**Figure 1B).** We identified numerous metabolic pathways that were distinctly increased in response to *G. vaginalis* as compared to *L. crispatus* exposure. These pathways appeared to be mostly consistent across cervicovaginal cell types. Those unique to *G. vaginalis* across all epithelial cell types notably included spermidine and spermine biosynthesis, as well as sphingolipid and purine/pyrimidine metabolism pathways. In contrast, glutamate metabolism, the Warburg effect, citric acid cycle and carnitine synthesis pathways were enriched in *L. crispatus*-exposed cervicovaginal cells.

Given the distinct alterations in spermidine and spermine biosynthesis, as well as observed differences in putrescine and spermidine metabolites in response to *G. vaginalis*, we next sought to investigate the impact of *G. vaginalis* and *L. crispatus*-exposures on the polyamine biosynthesis pathway specifically (**Figure 1C**). Spermidine levels were substantially elevated in *G. vaginalis*-exposed cells across all cell types. Ornithine and putrescine levels showed cell type-specific responses, with increased levels observed in Endo and VK2 cells exposed to *G. vaginalis*. Correlation analysis further revealed positive correlations between guanosine, cytidine 5’-monophosphate, and putrescine in Ecto cells; ornithine, pyruvate, and putrescine in Endo cells; and ornithine, guanosine, spermidine, and N’-acetylspermidine in VK2 cells (**Figure S3**). Taken together, these results identified metabolites that were altered across each cervicovaginal cell type, where impacts on putrescine and spermidine appear to be uniquely driven by *G. vaginalis* exposure.

### Exposure to polyamines shifts the *G. vaginalis*-mediated transcriptional response of Ecto, Endo and VK2 cells

Given the significant alterations in metabolomic profiles and the increase in polyamine biosynthesis in *G. vaginalis,* but not *L. crispatus*-exposed cells, we next sought to determine if exposure to polyamines would alter *G. vaginalis*-mediated changes in epithelial cell function. We hypothesized that epithelial cell production of polyamines may serve as a host defense mechanism against *G. vaginalis*, and that polyamine pretreatment may, therefore, attenuate microbe-induced epithelial dysfunction. To assess the global transcriptional impact of polyamines and *G. vaginalis*, we performed RNA-sequencing followed by principal component analysis (PCA) (**Figure 2A**). The first principal component (PC1) separated *G. vaginalis*- exposed samples from unexposed controls, while the second principal component (PC2) distinguished samples exposed to PUT regardless of *G. vaginalis* exposure (**Figure 2A**). Additional analysis revealed a positive correlation between PC3 and *G. vaginalis* exposure in VK2 cells, and PC1 and *G. vaginalis* exposure in Ecto and Endo cells (**Figure S4A**). We also found that treatment with polyamines was associated with PC2 across each cell type. Overall, these data suggest large shifts in the transcriptome in response to *G. vaginalis* treatment and that these shifts may be modified by treatment with polyamines.

To further explore the transcriptional effect of polyamines on *G. vaginalis*-mediated epithelial cell function, we performed differential expression (DE) analysis to identify DE genes (DEGs) in each polyamine+*G. vaginalis* (GV) condition compared to non-treated (NT)+GV. Additionally, we included the comparison between NT+GV and NT+noGV to determine the effects of *G. vaginalis* alone on transcriptomic signatures as a control for polyamine pretreatment. In most comparisons, there were hundreds of DEGs identified (**Figure 2B and Figure S4C**). The comparison between spermidine (SPD)+GV and NT+GV conditions, however, yielded few DEGs (9 genes for Ecto, 6 genes for Endo and 9 genes for VK2 cells). Volcano plots revealed polyamine alterations of *G. vaginalis*-induced genes known to be important to the host-defense response, such as epithelial barrier maintenance and inflammation, including CXCL8, CCL20, SP100A9, MMP9 and KRT13 (**Figure S4C**).

To identify if individual polyamines (PUT, SPM, SPD) upregulated unique genes during *G. vaginalis* exposure, we next generated UpSet plots (**Figure 2B**) of significant DEGs for each cell type exposed to polyamine+GV compared to NT+GV. Interestingly, exposure to *G. vaginalis* alone (NT+GV vs NT+noGV) led to the highest number of DEGs in Endo cells while pretreatment with SPM (SPM+GV) resulted in the highest number of altered genes in Ecto and VK2 cells. We also found that many genes were unique to each polyamine pretreatment in the presence of *G. vaginalis* across each cell type.

Functional annotation of these genes was performed using Metascape (**Figure 2C**). This analysis revealed that *G. vaginalis* exposure upregulated epithelial cell functional pathways related to bacterial defense (antimicrobial), cytokine stimulus/production, inflammatory response, and cell motility. Cell to cell adhesion pathways were upregulated after PUT+GV exposures in Endo and VK2 cells. In SPM+GV exposures, all three cell types exhibited upregulation of pathways related to bacterial defense responses. Interestingly, Ecto cells showed upregulation of pathways related to inflammatory responses and epigenetic processes after SPM+GV exposure. Overall, these data suggest that polyamines, and specifically treatment with putrescine, may alter the epithelial response to *G. vaginalis* exposure, transcriptionally driving cells towards a less inflammatory phenotype.

Lastly, we investigated whether *G. vaginalis* alters genes directly involved in polyamine biosynthesis and if pretreatment with polyamines themselves could modify these *G. vaginalis*- induced changes. To this end, we visualized the expression of genes encoding enzymes involved in polyamine synthesis and catabolism (**Figure 2D**). Our analysis revealed distinct cell type-specific responses in *G. vaginalis*-exposed epithelial cells with putrescine (PUT+GV) or spermine (SPM+GV) pretreatment. Notably, SAT1 (spermidine/spermine N(1)-acetyltransferase 1), an enzyme responsible for spermine and spermidine catabolism, was universally upregulated across all three cervicovaginal cell types in response to SPM+GV suggesting a negative feedback loop that may prevent overproduction of spermine and spermidine. A similar effect was seen with an upregulation of PAOX (Peroxisomal N(1)-acetyl-spermine/spermidine oxidase), a catabolism enzyme that oxidizes N1-acetylspermine to spermine and N1-acetylspermidine to spermidine, after exposure to SPM+GV in VK2 cells. AMD1 (adenosylmethionine decarboxylase 1), an enzyme that catalyzes the biosynthesis of spermine and spermidine from putrescine, was also upregulated in all three cervicovaginal epithelial cell types after PUT+GV exposure providing evidence that putrescine increases polyamine biosynthesis. Overall, these results suggest that polyamine pretreatment may lead to positive feedback of polyamine biosynthesis that increases the protective impact of polyamines on cervicovaginal epithelial cells.

### WGCNA reveals de novo gene modules associated with polyamine exposure in the presence of *G. vaginalis*

Since pretreatment with polyamines showed significant impacts on *G. vaginalis*-induced transcriptional pathways, we next wanted to identify de novo modules of genes that were associated with these polyamine effects. Given limitations of DEG functional annotation and the inability to determine particular genes of interest, WCGNA analysis was used as a secondary approach to examine gene modules altered by polyamines in the context of *G. vaginalis* exposure. We focused this analysis on *G. vaginalis* exposed epithelial cells that were pretreated with putrescine or spermine since so few genes were altered with spermidine pretreatment. We performed WGCNA on transcriptomic data across each treatment and cell type. We identified nine total modules (**Figure 3A**). Overall, hierarchical clustering of these modules was associated with cell type. The module green-yellow correlated with Endo cells, the salmon module with VK2 cells, and the dark orange module with Ecto cells regardless of polyamine or bacterial exposure conditions (**Figure 3A, Figure S5**). After running pathway analysis on each of these cell type modules, we found that enriched functions (**Table S5**) included typical cellular processes (metabolism of lipids and fatty acids) as well as cellular response to cytokines for Ecto cells, cell cycle regulation and chromosome organization for Endo cells and extracellular matrix organization, cellular adhesion and leukocyte migration in vaginal cells.

We next wanted to determine which modules drive the functional effects of polyamines in the setting of *G. vaginalis* exposure. To obtain co-expression signatures indicating differences in gene alterations with or without polyamine pretreatment in *G. vaginalis* exposed cells, we merged the polyamine and *G. vaginalis* exposure conditions across the cell types and performed module eigengene analysis (**Figure S5**). Through this analysis, we found that the dark green module included a group of genes that were not transcriptionally altered by *G. vaginalis* exposure alone but were changed in the presence of polyamines and *G. vaginalis*. By using this module, we were able to focus on genes that are specifically altered by polyamines only when *G. vaginalis* is present (**Figure 3B**). This module was enriched for several functions related to epithelial cell differentiation, an interconnected network of NF-kB activation and signaling, and a small group of functions related to SUMOylation (**Figure 3C**).

To better specify the genes in this module and their functions, we next performed a merged network analysis. A co-expression network of genes in the dark green module was built by obtaining nodes and edges separately for each polyamine condition. This network was then visualized for the top 50 genes associated with PUT+GV treatment across all three cell types (**Figure 3C**), with the top 10 genes highlighted in yellow based on significant fold change relative to non-treated controls (NT+GV) (**Figure 3D**). The resulting network revealed a dense and interconnected set of genes that are altered by putrescine pretreatment in the context of *G. vaginalis* exposure. These include PCDH10, a protocadherin gene involved in cell adhesion and epithelial to mesenchymal transition (EMT), and ID2 which is involved in cell proliferation. We performed a similar analysis to investigate the effect of SPM on *G. vaginalis*-mediated gene transcription and found a similar highly connected network of genes (**Figure 3F**) with SNAl1, a prominent EMT gene, being highly connected to multiple other genes. Many of the top 10 genes in this network also had elevated fold changes relative to non-treated controls (NT+GV) (**Figure 3G**). Overall, we identified a module of genes (module dark green) that is associated with the impacts of putrescine or spermine pretreatment on *G. vaginalis* exposure, suggesting several core epithelial cell functions that may be altered including those related to epithelial barrier integrity and inflammation.

### Exposure to *G. vaginalis* in the presence of polyamines leads to attenuated inflammatory responses that vary across cell types

Since pretreatment with polyamines altered the global transcriptome of *G. vaginalis* exposed cervicovaginal epithelial cells, including factors related to inflammation and epithelial cell differentiation, we next determined if these effects could be detected in the secreted proteome. We performed Luminex analysis on culture supernatants from Ecto, Endo and VK2 cells exposed to *G. vaginalis* with or without polyamine pretreatment. With analytes passing stringent quality control cutoffs, we performed PCA to determine if there were global shifts in our data. PC1 clearly separated *G. vaginalis*-exposed samples from non-treated controls (**Figure 4A**) across all three cell types, accounting for >80% of variability. We found a small separation along PC2 according to polyamine pretreatment, with some dispersion along PC2 in response to polyamines in the *G. vaginalis*-exposed conditions.

**Figure 4.**
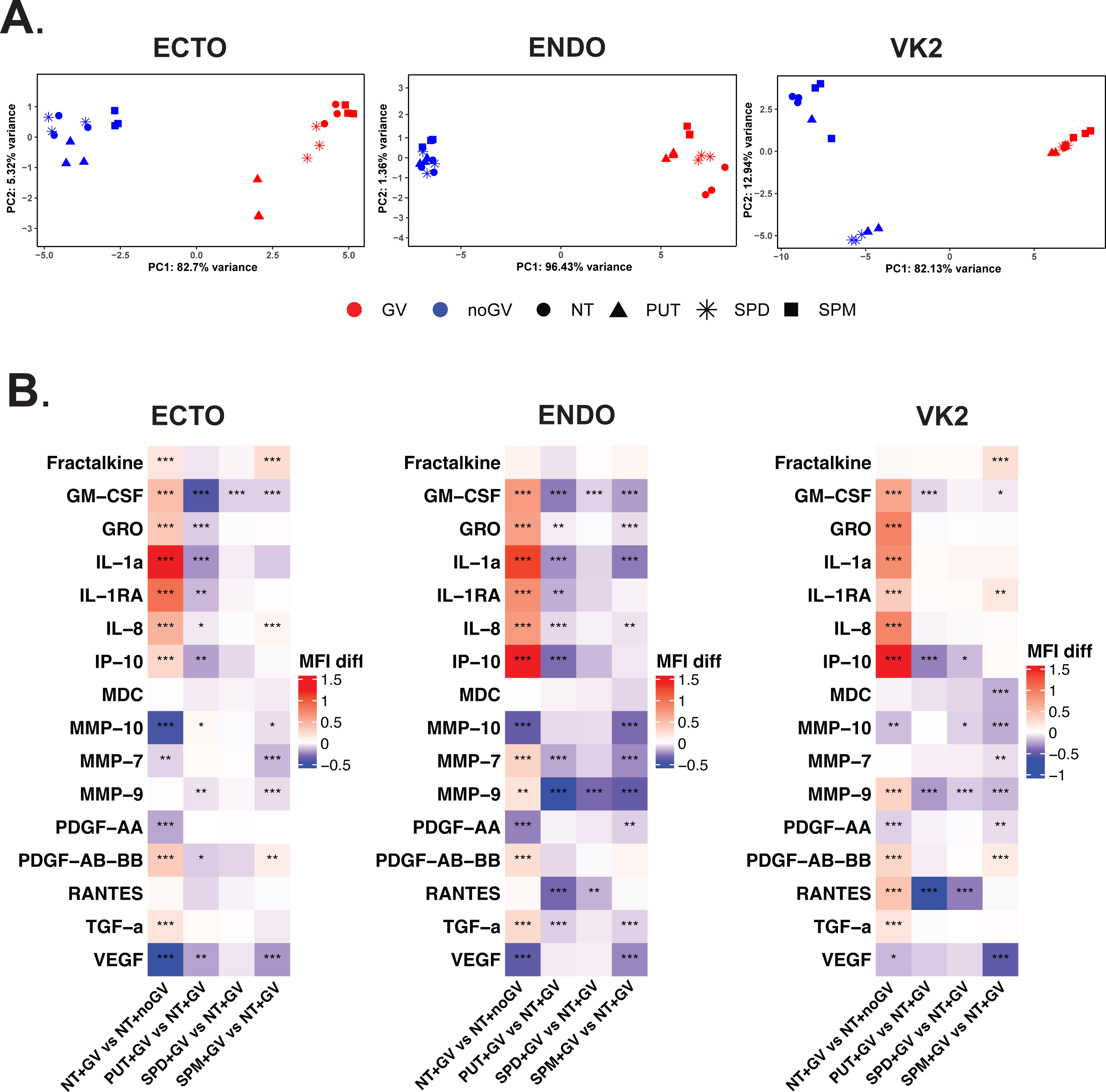
Polyamines modify *G. vaginalis-*induced inflammatory response. (A) PCA plots along the first two principal components shown for Ecto, Endo, and VK2 lines by polyamine pretreatment (NT, PUT, SPD, or SPM) and colored by GV (red) or no GV (blue) exposure. (B) Heatmap representing results of two-way ANOVA test and Tukey’s Honest Significant Difference (TukeyHSD) post-hoc test. Heatmap corresponds to MFI difference (MFI diff) between combined levels of treatment and GV exposure (e.g., PUT+GV vs NT+GV). Asterisks signify significance by p-value; *p<0.05, **0.05<p<0.01, ***p<0.01.

The MFI signals were visualized in a heatmap with corresponding model results indicated in **Figure 4B**. Notably, exposure to *G. vaginalis* (NT+GV vs NT+noGV) significantly altered cytokine concentrations, markers of an activated inflammatory response, in Ecto (nine increased, two decreased), Endo (eight increased, two decreased) and VK2 (nine increased, two decreased) cells. Putrescine pretreatment (PUT+GV vs NT+GV) significantly decreased the *G. vaginalis*-induced proinflammatory cytokines in Ecto (eight cytokines), Endo (eight cytokines) and VK2 (three cytokines) cells. Spermine pretreatment (SPM+GV vs NT+GV) also significantly reduced *G. vaginalis*-induced cytokines in Ecto (two cytokines), Endo (seven cytokines) and VK2 (four cytokines) cells. In addition to cytokines, a panel of matrix metalloproteinases (MMPs), enzymes known to cleave tight junctions and increase epithelial barrier permeability, were measured by Luminex. *G. vaginalis* (NT+GV vs NT+noGV) altered MMP concentrations in Ecto (two decreased), Endo (two increased, one decreased) and VK2 (one increased, one decreased) cells. *G. vaginalis*-induced MMPs including MMP-7 and MMP-9 were decreased with putrescine (PUT+GV vs NT+GV) and spermine (SPM+GV vs NT+GV) pretreatment in Endo and VK2 cells. While spermidine pretreatment (SPD+GV vs NT+GV) also reduced cytokine and MMP concentrations, similar to transcription results, there were few that reached significance in Ecto (one cytokine), Endo (two cytokines, one MMP) and VK2 (two cytokines, two MMPs) cells. These results provide evidence of a protective functional effect of polyamines, specifically putrescine and spermine, as they mitigate *G. vaginalis*-induced host inflammatory responses by decreasing pro-inflammatory cytokines and reducing tight junction cleavage through the mitigation of MMP secretion.

### Correlation of *in vitro* transcriptional signatures to existing human datasets

We next investigated if our *in vitro* findings, showing that polyamines mitigate *G. vaginalis*-induced transcriptomic changes, could be validated in an *in vivo* setting. To do this, we leveraged existing transcriptomic data from clinical studies on BV and preterm birth, including Wikstrom et al. (preterm birth signature)^61^ and Berard et al. (*Lactobacillus*-dominant signature)^24^ (**Table 1)**. To determine if there were significant overlaps between our gene profiles and the gene signatures reported in these clinical studies, we compared the upregulated genes in *G. vaginalis*- exposed cervicovaginal epithelial cells either alone or with putrescine or spermine pretreatment to differentially expressed genes reported in the human studies. The resulting heatmap highlights significant gene overlaps in cells exposed to *G. vaginalis* with polyamine pretreatment (PUT+GV vs NT+GV or SPM+GV vs NT+GV) with signatures of a *Lactobacillus*-dominant microbiome. (**Figure 5A, Table S6**). As expected, significant overlap in gene signatures were also found between preterm birth and *G. vaginalis* exposed vaginal cells (NT+GV vs NT+noGV). However, Ecto and Endo cells exposed to *G. vaginalis*, while still showing an increased trend for overlapping genes with the preterm birth cohort, did not reach statistical significance. Interestingly, in the presence of polyamines (PUT+GV vs NT+GV or SPM+GV vs NT+GV), the overlap of genes with preterm birth is decreased in all three cell types. Taken together, these findings suggest that polyamines, and specifically putrescine and spermine, may shift epithelial transcriptomic responses towards those that are more like a *Lactobacillus*- dominant vaginal microbiome or less like those seen in a preterm birth.

**Figure 5.**
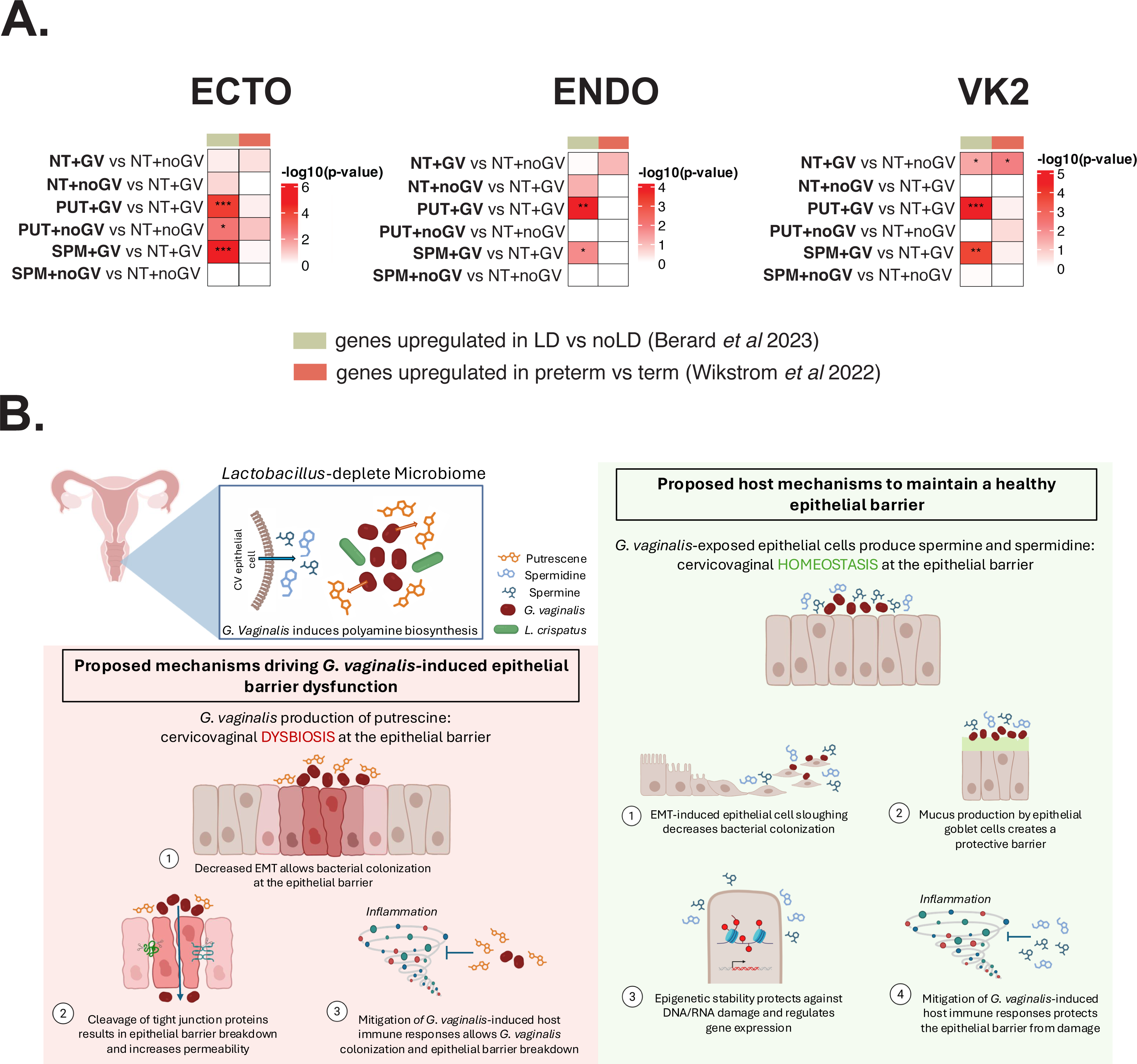
Comparison of *in vitro* transcriptional signatures to existing human datasets. (A) Heatmap summarizes output of Fisher Exact Test showing significant overlap in upregulated genes in each in vitro comparison (our analysis) and in vivo clinical datasets. Significance marked with asterisks; *p<0.05, **p<0.01, ***p<0.001 (B) Graphical schematic representing study findings and proposed mechanisms of how polyamines alter *G. vaginalis*-induced epithelial dysfunction and their contribution to cervicovaginal health and disease.

**Table 1.** Description of human clinical studies used for correlation to *in vitro* transcriptomic signatures.

| Study | Study Type | Study Location | Subject Info | Sample Type | Outcome | Transcriptomics Assay | *In vitro* comparison to: |
| --- | --- | --- | --- | --- | --- | --- | --- |
| Berard, et. al<sup>1</sup> | Nested case-control Partners PrEP Study | Kenya and Uganda | Non-pregnant: 25-51 years of age | Vaginal biopsies (n=80) | Microbiome -associated epithelial dysfunction | Illumina Beadchip gene expression microarray | Transcriptomics results from *Lactobacillus*-dominant (LD) vs non-*Lactobacillus*-dominant (noLD) microbiome |
| Wikstrom, et. al<sup>2</sup> | Nested case-control CERVIX Study | Sweden | Pregnant: 18-20.6 weeks of gestation | Cervical swabs (n=144) | Preterm Birth | Illumina TrueSeq Total Strand RNA, NovaSeq 6000 | Transcriptomics results from preterm vs term deliveries |

1. Berard, A.R., Brubaker, D.K., Birse, K., Lamont, A., Mackelprang, R.D., Noël-Romas, L., Perner, M., Hou, X., Irungu, E., Mugo, N., et al. (2023). Vaginal epithelial dysfunction is mediated by the microbiome, metabolome, and mTOR signaling. *Cell Rep* 42, 112474. 10.1016/j.celrep.2023.112474.
2. Wikström, T., Abrahamsson, S., Bengtsson-Palme, J., Ek, J., Kuusela, P., Rekabdar, E., Lindgren, P., Wennerholm, U.B., Jacobsson, B., Valentin, L., and Hagberg, H. (2022). Microbial and human transcriptome in vaginal fluid at midgestation: Association with spontaneous preterm delivery. *Clin Transl Med* 12, e1023. 10.1002/ctm2.1023.

## Discussion

This study shows microbe-specific alterations in global metabolite production by bacteria-exposed cervicovaginal epithelial cells. Metabolite profiles suggest that *G. vaginalis* promotes polyamine biosynthesis. Vaginal polyamines alter *G. vaginalis*-mediated epithelial cell transcription, signifying functional changes in host immune responses, epithelial cell differentiation, and epigenetic processes. Transcriptomic changes in enzymes responsible for polyamine synthesis and catabolism support our metabolomic findings and reveal a potential mechanism by which *G. vaginalis* drives alterations in polyamine biogenesis in the cervicovaginal epithelium. Additionally, polyamines constrain *G. vaginalis*-induced secretion of proinflammatory cytokines and matrix metalloproteinases, suggesting that vaginal polyamines can regulate epithelial immune responses to vaginal microbes. These results highlight the complexity of the cervicovaginal ecosystem, in which interactions between the microbiome and microbial/host-derived metabolites play critical roles in regulating epithelial cell function and host immune responses.

Elucidating host-microbe interactions in the cervicovaginal space are critical to identifying the molecular mechanisms linking epithelial dysfunction to adverse reproductive outcomes. In this study, exposure of cervicovaginal epithelial cells to microbes representing either a *Lactobacillus*-dominant (*L. crispatus*) or a *Lactobacillus*-deplete (*G. vaginalis*) microbiome revealed strain-specific alterations of the metabolome. Unsurprisingly, in cervicovaginal epithelial cells exposed to *L. crispatus*, a microbe associated with vaginal health, we detected increased secretion of lactic acid derived metabolites, including imidazole lactate, phenylactate and indolelactate. These lactate derivatives contribute to the vaginal acidic pH associated with the protective properties of a *Lactobacillus*-dominant microbiome.^10,62^ Interestingly, the Warburg effect, used to describe the increase of lactate production in cancer cells,^63^ was identified as an increased pathway in *L. crispatus* exposed cells in our metabolomic results. Succinate was also consistently induced by *L. crispatus* exposure in all three cervicovaginal cell types. Despite succinate being previously associated with BV and proinflammatory or immunomodulatory effects,^64–66^ our study and others have shown that *L. crispatus*-dominate microbiomes also produce characteristically high succinate levels.^67,68^ In an animal model, succinate has also been reported to have protective effects in the intestine, including expression of tight junction proteins (claudin and occludins) that tighten the epithelial barrier and protect against ischemia reperfusion injury.^69^ The contradictory results of these studies add to the complexity of elucidating metabolite contributions to epithelial cell function without investigation into other parts of the cervicovaginal ecosystem including host-defense mechanisms.

Oppositely, cervicovaginal cells exposed to *G. vaginalis*, a microbe predominately associated with BV and adverse obstetrical outcomes,^4,15–17,19^ altered the production of multiple metabolites belonging to the polyamine biosynthesis pathway. Altered polyamine metabolites included an increase in putrescine, spermidine, N-acetylglutamate (the first step in arginine biosynthesis and catalyst for polyamine biosynthesis), and decreased 5- methylthioadenosine (MTA), a breakdown product of the conversion of putrescine to spermidine and spermidine to spermine. Putrescine is often found in high concentrations in vaginal fluid, is associated with malodor in clinical cases of BV characterized by a *Lactobacillus*-deplete microbiome and is believed to be a clinical biomarker for BV diagnosis.^64,70^ However, the contribution of *G. vaginalis* to these findings has been unclear, as several studies investigating the vaginal metabolome in non-pregnant women with BV have not been able to find direct evidence linking *G. vaginalis* to increased putrescine production and instead have found associations with other BV-associated bacteria.^45,71^ The discrepancy between these previous studies and our observation of increased putrescine after *G. vaginalis* exposure may be due to the strain of *G. vaginalis* that was studied. As recent studies have identified many different *G. vaginalis* clades with strain-specific virulence factors,^72–74^ the metabolites produced might differ significantly, making characterization of *G. vaginalis* as a singular bacteria more complex. Interestingly, we also saw an induction of spermidine in all three cervicovaginal epithelial cell types, which has not been previously associated with *G. vaginalis*. However, as spermine and spermidine often originate from eukaryotic cells,^75,76^ it is more likely that increased spermidine production is a cervicovaginal epithelial cell defense response to protect against the effects of *G. vaginalis*. Whether this specific strain of *G. vaginalis* has the enzymes necessary to produce spermine is unknown. These results also suggest that metabolic regulation of host-defense could play a key role in protection of the cervicovaginal epithelial barrier during *G. vaginalis* colonization.

The metabolomics results from this study suggest a role for *G. vaginalis* in regulating polyamine biosynthesis which consequently could have functional effects on cervicovaginal epithelial cells. Follow up transcriptomics revealed significant functional alterations in all three cervicovaginal cell types with polyamine treatment prior to *G. vaginalis* exposure. As has been shown previously,^33^ *G. vaginalis* significantly altered transcription of genes and functional pathways related to host-defense, innate immune response, apoptosis and cellular adhesion when compared to non-treated cells. These effects of *G. vaginalis* on cervicovaginal epithelial cell function have been shown previously by our group and others.^20,29–31,77^ More interestingly, pretreatment with polyamines, especially putrescine and spermine, prior to *G. vaginalis* exposure resulted in significant transcriptional shifts not seen with polyamine treatment alone, indicating that polyamine-induced alterations in epithelial cell function are dependent upon the cervicovaginal microbiome. As there was very little overlap in altered genes between polyamine exposures, the transcription changes were largely polyamine specific, providing evidence of a regulated effect on *G. vaginalis*-induced cellular function. Surprisingly, both putrescine and spermine mitigated the *G. vaginalis*-mediated transcription of genes related to an inflammatory response, cytokine production and bacterial defense, suggesting that both polyamines can reduce the epithelial host-defense response initiated by exposure to *G. vaginalis*. This was further evidenced by the reduction in *G. vaginalis*-induced proinflammatory cytokines, chemokines and matrix metalloproteinases (e.g., IL-8, GM-CSF, IL-1α, and MMP9 among others) by putrescine and spermine. While these results indicate that putrescine and spermine both mitigate cervicovaginal inflammation, it is plausible that putrescine produced predominately by anaerobes such as *G. vaginalis* during BV infection, allows *G. vaginalis* to evade the host immune response by attenuating cytokine production. Conversely, spermine, largely produced by eukaryotic cells, acts as a part of the host defense response to protect the cervicovaginal epithelial barrier from the negative effects of *G. vaginalis* colonization. This highlights the complexity of the metabolome when metabolites from the same biosynthesis pathway exert similar molecular effects but drive opposing physiological endpoints.

Also of particular interest, spermine pretreatment prior to *G. vaginalis* exposure activated gene pathways associated with epigenetic processes across the three cervicovaginal epithelial cell types. In eukaryotic cells, polyamines are known to predominantly bind RNA and DNA due to their net positive charge at a physiological pH and stabilize RNA and DNA structure.^75,78^ It is therefore feasible that a *Lactobacillus*-deplete microbiome, with a less acidic pH, could result in epithelial cell host-derived increases in spermine that binds to DNA structures resulting in epigenomic stabilization. Interestingly, *L. crispatus* has been shown to use similar epigenetic mechanisms to decrease chromatin accessibility as a possible mechanism to protect against epigenomic damage by microbial pathogens and to suppress the immune sytem.^33,79,80^ While our understanding of epigenetic regulation by vaginal bacteria is in its early stages, studies have shown that bacterial production of lactate can increase histone lactylation in macrophages promoting cellular homeostasis.^81^ *L. crispatus* has been shown to decrease *Chlamydia trachomatis* infection in cervicovaginal epithelial cells by decreasing histone deacetylase HDAC4 and increasing histone acetylase (HAT) EP300 through the production of D(-) lactic acid.^79,82^ Histone acetylation alters DNA-histone binding and allows the transcription of genes leading to changes in epithelial cell function including cell proliferation which was a focus of these studies.^79,82,83^ Future studies elucidating the role of lactate metabolites in the regulation of epigenetic modifications in cervicovaginal epithelial cells would be critical to the development of lactate-focused therapies to prevent adverse gynecological and reproductive adverse outcomes.

Further investigation into de novo gene modules showing large differences in gene expression between *G. vaginalis* exposure with or without polyamine pretreatment identified interconnected gene networks regulating epithelial function. When cells were pretreated with putrescine prior to *G. vaginalis* exposure, genes known to increase cell proliferation (ID2, ACTBL2)^84^, decrease epithelial-mesenchymal transition (PCDH10, ID2)^85,86^ and reduce cellular adhesion (PCDH10)^87^ were activated. These results suggest that putrescine may prevent EMT-mediated epithelial cell shedding described as a defense response against microbial pathogen colonization. During Group B Streptococcus (GBS) infection, epithelial cell sloughing resulting from EMT, is increased as part of the host defense process to prevent pathogenic bacteria from gaining access to the epithelial cell barrier.^88^ Additionally, *G. vaginalis* has been shown to cleave epithelial cell tight junctions as a mechanism to break down the cervicovaginal barrier.^21,30,89^ These transcriptional changes suggest that putrescine may function as a *G. vaginalis* effector molecule, promoting bacterial colonization and compromising integrity of the epithelial barrier. On the other hand, cervicovaginal cell exposure to spermine prior to *G. vaginalis* resulted in a unique set of altered genes that targeted many of the same functions as those altered by putrescine but with differential biological outcomes. Spermine upregulated genes that promote epithelial-mesenchymal transition (SNAI1, NEDD9),^90,91^ cell proliferation and migration (NEDD9)^91^ possibly in an attempt to increase epithelial cell shedding to prevent colonization of bacteria at epithelial barriers. Additionally, spermine increased genes with very specialized functions such as DLL1, which regulates epithelial cell fate during stem cell differentiation and has been shown to cause regeneration of mucus-producing epithelial goblet cells.^92^ As the mucosal barrier is a critical part of the host-defense response to invading pathogens, increased mucus production could be a compensatory epithelial cell response to protect the epithelial barrier. Overall, these results provide evidence that polyamines regulate multifaceted and complex epithelial cell functions after *G. vaginalis* colonization by mediating inflammatory host-defense responses, epithelial barrier integrity, epigenetic protection and mucus production.

While the results of this study provide evidence that polyamines can alter *G. vaginalis*- induced transcriptional changes in cervicovaginal epithelial cells, we wanted to determine if these *in vitro* results correlated with findings from human studies. Comparisons of transcriptional data sets from two studies focused a *Lactobacillus*-deplete microbiome^24^ or preterm birth^61^ showed partial overlap in gene signatures with our epithelial cell findings. Perhaps expectedly, there is a significant overlap in gene signatures between *G. vaginalis*-exposed vaginal cells and those seen with preterm birth in the Wikstrom, et. al study. A similar, although non-significant trend, was seen in cervical cells. Interestingly, this overlap appears to decrease with polyamine exposure across all three cell types suggesting that polyamines may be able to shift transcriptomic signatures away from those associated with preterm birth. When comparing the transcriptomic results of the spermine pretreated *G. vaginalis*-exposed cells to those of the *Lactobacillus*-dominant samples in the Berard, et. al study, there is a significant overlap between the transcriptional profiles in both studies in all three cervicovaginal epithelial cell types. These results suggest that spermine can create a gene transcription signature that is similar to a *Lactobacillus*-dominant microbiome, despite the presence of *G. vaginalis*. This would further support our findings that spermine, in the presence of *G. vaginalis*, is able to provide protection against *G. vaginalis*-induced changes in cervicovaginal epithelial cell function. Perhaps unexpectedly, we saw the same response with putrescine pretreatment. However, considering that our transcriptomic and Luminex results suggested that putrescine reduced the host bacterial defense and inflammatory response in *G. vaginalis*-induced epithelial cells, it would not be as surprising to see an association with transcriptomic results from a *Lactobacillus*-dominant microbiome. These results provide greater emphasis on the need for increased investigation into functional assays to assess the role of transcriptional changes on epithelial cell function.

Nonetheless, comparisons of transcriptional signatures between *in vitro* and relevant *in vivo* studies provide evidence for the validity of our findings in cervicovaginal epithelial cells with polyamines acting to protect against cellular dysfunction and adverse clinical outcomes.

We acknowledge several limitations to this study, namely the use of singular bacteria species. While *L. crispatus* and *G. vaginalis* were chosen as common representative strains of either a *Lactobacillus*-deplete or *Lactobacillus*-dominant microbiome, there are other prominent CST IV bacterial strains (*Prevotella* ssp., *Snethia* ssp., *Atopobium* spp., etc.) that need to be explored. Additionally, multi-species microbial communities representing CST I or CST IV should be further studied to identify the effects of microbe-microbe interactions on metabolome and transcriptional endpoints. However, it is critical to first elucidate the molecular effects of singular microbial species to more clearly identify mechanistic areas of interest before moving into investigations with larger more complex multi-species communities. Additionally, this study does not address the contribution of cervicovaginal immune cells to the metabolome or how their function may be altered by polyamines. This work would require further follow-up studies investigating the effects of polyamines on epithelial-immune cell communication.

Overall, this study provides insight into the effects of microbiome-metabolome interactions on epithelial cell function in the cervicovaginal space. The results of this study identify specific metabolites contributing to the *G. vaginalis*-mediated alterations in epithelial cell function. The induction of polyamine biosynthesis by *G. vaginalis*, but not *L. crispatus*, provides evidence of a microbe-specific role of polyamines in regulating epithelial cell host-defense responses. These findings begin to ascribe critical mechanistic functions to metabolites in the cervicovaginal space. Future studies focused on understanding the role of putrescine and spermine in epithelial immune regulation and barrier integrity will be essential to the development of potential new postbiotic therapies to prevent microbe-induced adverse gynecological and obstetrical outcomes.

## Author contributions

Conceptualization: KDG, LA

Experiments and Data Collection: KDG, LA, BF, KK

Data analysis, bioinformatics, and modeling: BAG, OK, RP

Paper writing and first draft: LA, BAG, KDG, OK, RP

Figure generation: OK, RP, BAG, LA, KDG

Draft reviewing and editing: LA, BAG, KDG, OK

Supervision: KDG, BAG, LA

Funding: BAG, KDG

## Acknowledgements

The authors would like to acknowledge Dr. Elliot Friedman and Dylan Curry in the Microbial Culture and Metabolomics Core of the PennCHOP Microbiome Program for providing microbial culture services for these studies. Additionally, the authors would like to thank Dr. Lynn Spruce in the High Throughput Sequencing Core at the Children’s Hospital of Pennsylvania (CHOP) for completing the RNA sequencing, as well as RNA-sequencing bioinformatics performed by John Tobias. We would also acknowledge Drs. Yangzhu Du, Honghong Sun, and Eline Luning Prak of the Human Immunology Core at the Perelman School of Medicine at the University of Pennsylvania for assistance with Luminex assay. The HIC is supported in part by NIH P30 AI045008 and P30 CA016520. HIC RRID: SCR_022380

## Supplemental Figure Captions

**Figure S1. Polyamine dose response reduces *G. vaginalis*-induced IL-8 in Endocervical cells.** Endocervical cells were used as a representative cell type when determining appropriate doses of polyamines for future experiments. Values are mean ± SEM. Asterisks over lines represent comparisons between treatment groups. Comparisons were made between non-treated (NT) and polyamines alone (for all doses) or GV vs polyamine+GV (for all doses). One-way ANOVA with Tukey’s post-hoc test for multiple comparisons. *p<0.05, **p<0.01, ***p<0.001, ****p<0.0001.

**Figure S2. Lactate Dehydrogenase (LDH) is unchanged with polyamine pretreatment in cervicovaginal epithelial cells.** LDH is released from the cell when the integrity of the cell membrane is disrupted and therefore is representative of cell death. *G. vaginalis* increased LDH in ectocervical, endocervical and vaginal epithelial cells. Values are mean ± SEM. Asterisks over lines represent comparisons between treatment groups. Comparisons were made between non-treated (NT) and polyamines alone or GV vs polyamine+GV. One-way ANOVA with Tukey’s post-hoc test for multiple comparisons. *p<0.05, **p<0.01, ***p<0.001, ****p<0.0001.

**Figure S3. Metabolites that correlate with putrescine, spermidine and N(‘1)- acetylspermidine in Ectocervical (Ecto), Endocervical (Endo) and Vaginal (VK2) cells.** (**A**) Venn diagrams of upregulated metabolites in GV vs NT (left) and LC vs NT exposures by cell type (right). (**B**) Bar plots for top 25 compounds that correlate with putrescine in Ecto (left), Endo (center) and VK2 (right) cell lines. (**C**) Bar plots for top 25 compounds that correlate with spermidine in Ecto (left), Endo (center) and VK2 (right) cell lines. (**D**) Bar plots for top 25 compounds that correlate with N(‘1)-acetylspermidine in Ecto (left), Endo (center) and VK2 (right) cell types.

**Figure S4. Additional transcriptomic analyses for Ectocervical (Ecto), Endocervical (Endo) and Vaginal (VK2) cells.** (**A**) Correlation plots between 20 principal components and metadata features in Ecto, Endo and VK2 cells. (**B**) Upset plots show overlapping genes for each indicated comparison by upregulated and downregulated gene lists, where input gene lists were filtered based on significance (padj < 0.05) and fold change (FC > 2). (**C**) Volcano plots for pairwise comparisons of interest in Ecto, Endo and VK2 cells. Upregulated genes highlighted in red and downregulated genes highlighted in blue. The top 10 DE genes were labeled.

**Figure S5. Violin plots show the relationship between module eigengene and bacterial exposure and polyamine pretreatment phenotypes in each cell type.** Eigengene values of representative WGCNA modules in (**A**) ectocervical (Ecto), (**B**) endocervical (Endo), and (**C**) vaginal (VK2) cell lines. Each boxplot shows module eigengene expression across polyamine treatments (none, putrescine, spermidine, spermine) and bacterial exposure statuses (GV, noGV). Statistical comparisons between bacterial exposure groups were performed using Student’s *t*-test; *p<0.05, **p<0.01, ***p < 0.001, ns = not significant.

**Supplemental Table 1**. Full Metabolomics results and associated metadata.

**Supplemental Table 2**. Full Luminex results.

**Supplemental Table 3.** Full two-way ANOVA results for Luminex.

**Supplemental Table 4.** Overlapping metabolites between cervicovaginal cell types.

**Supplemental Table 5.** GO Gene enrichment results from the WCGNA modules.

**Supplemental Table 6.** List of overlapping genes between *in vitro* and *in vivo* studies.

