## Supplementary figures and images for "Multiomic analysis reveals that polyamines alter *G. vaginalis*-induced cervicovaginal epithelial cell dysfunction"

### Supplemental Figure 1

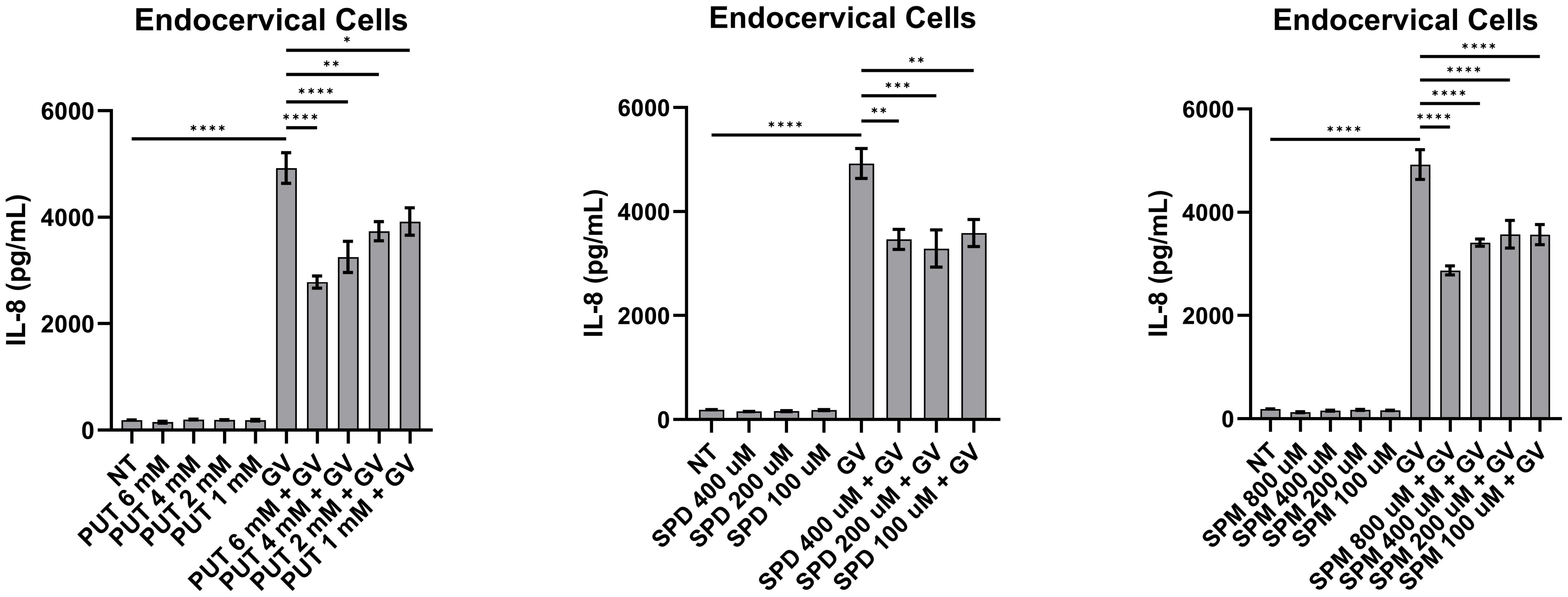

### Supplemental Figure 2

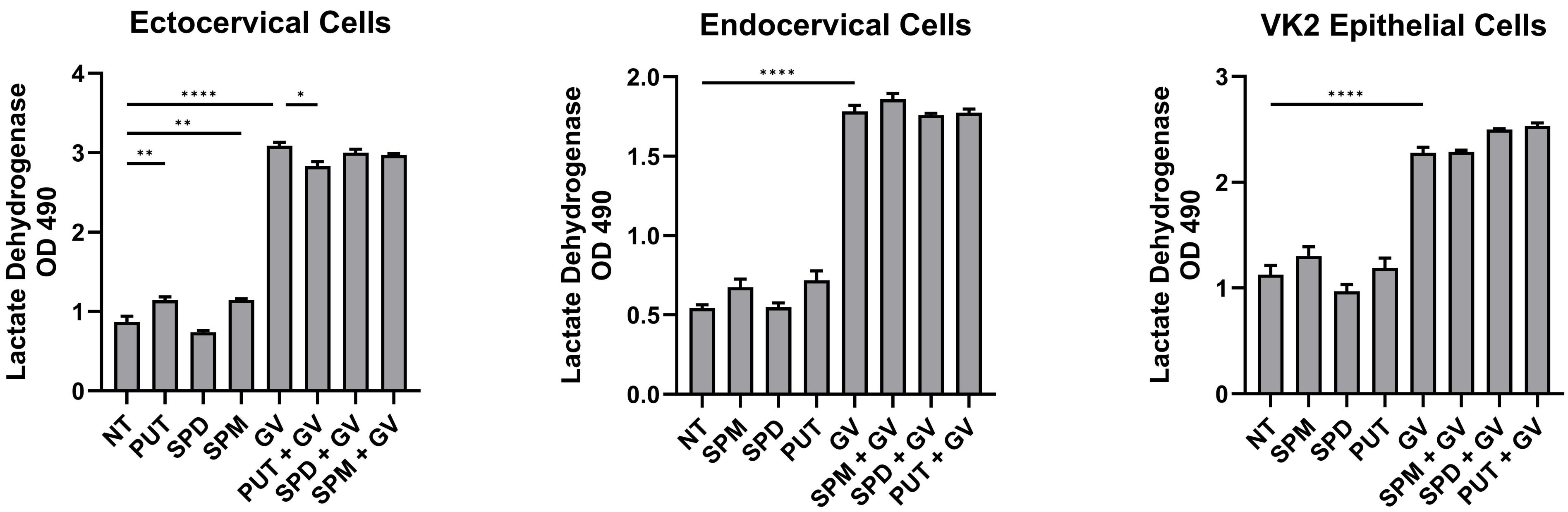

### Supplemental Figure 3

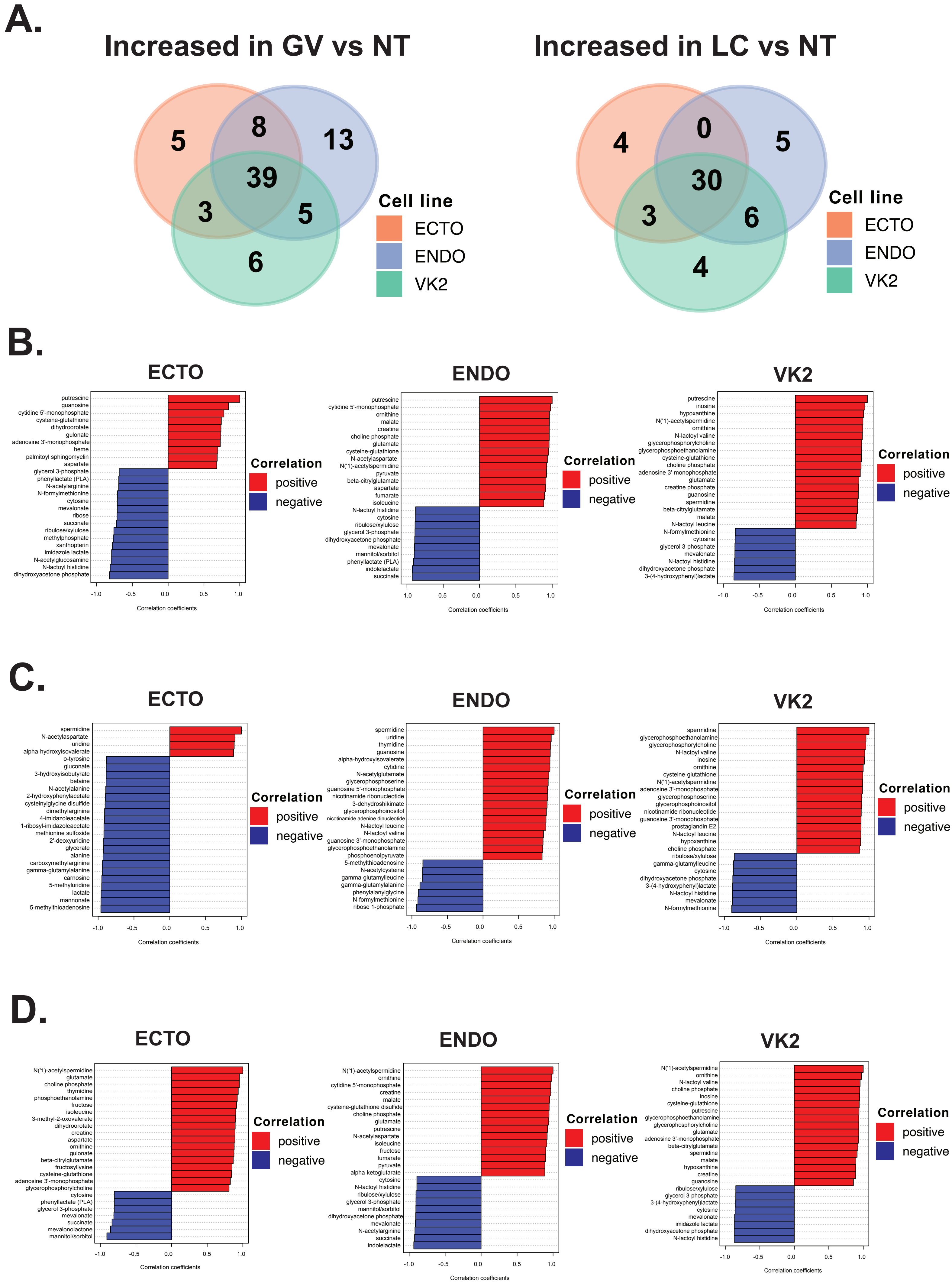

### Supplemental Figure 4

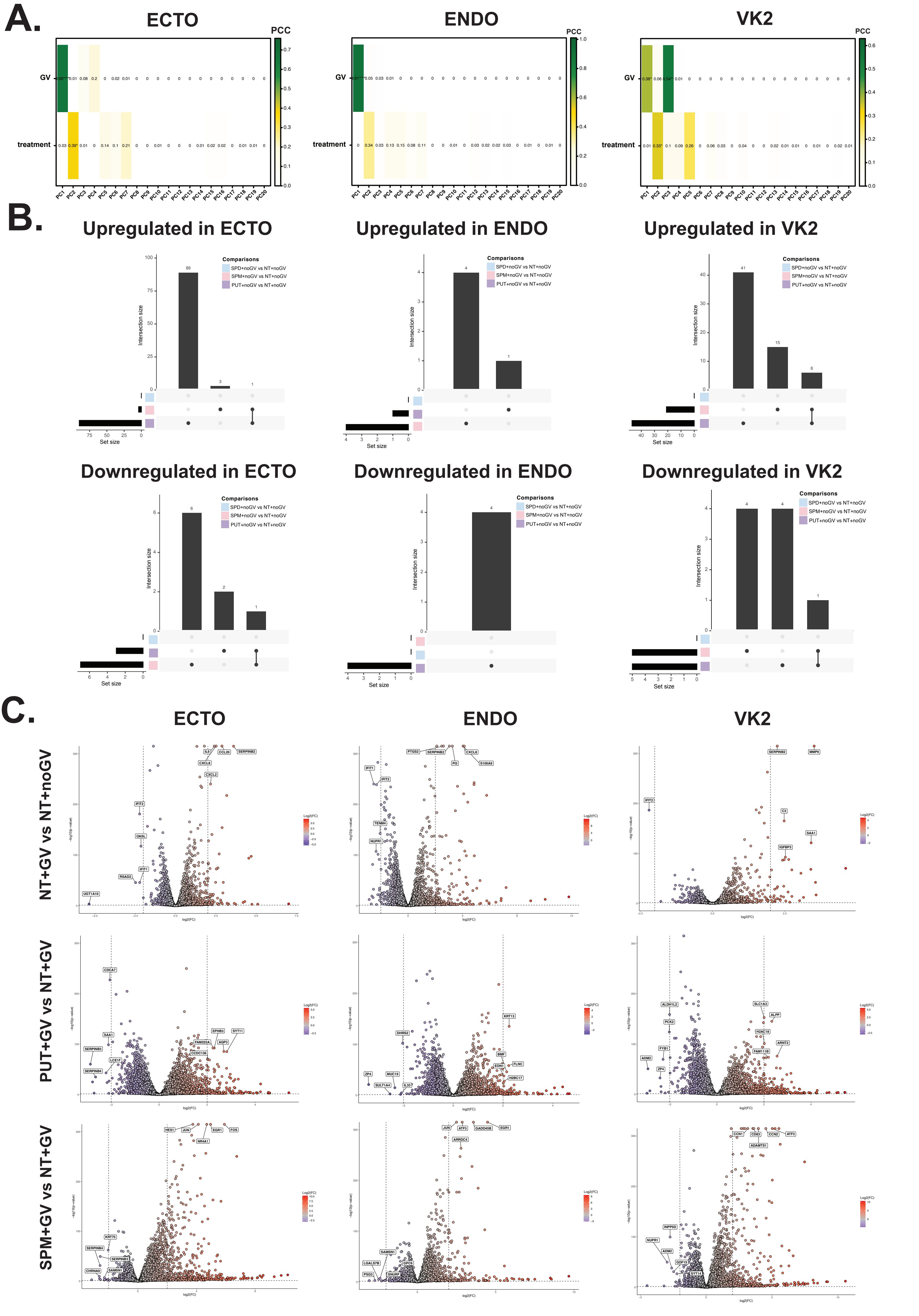

### Supplemental Figure 5

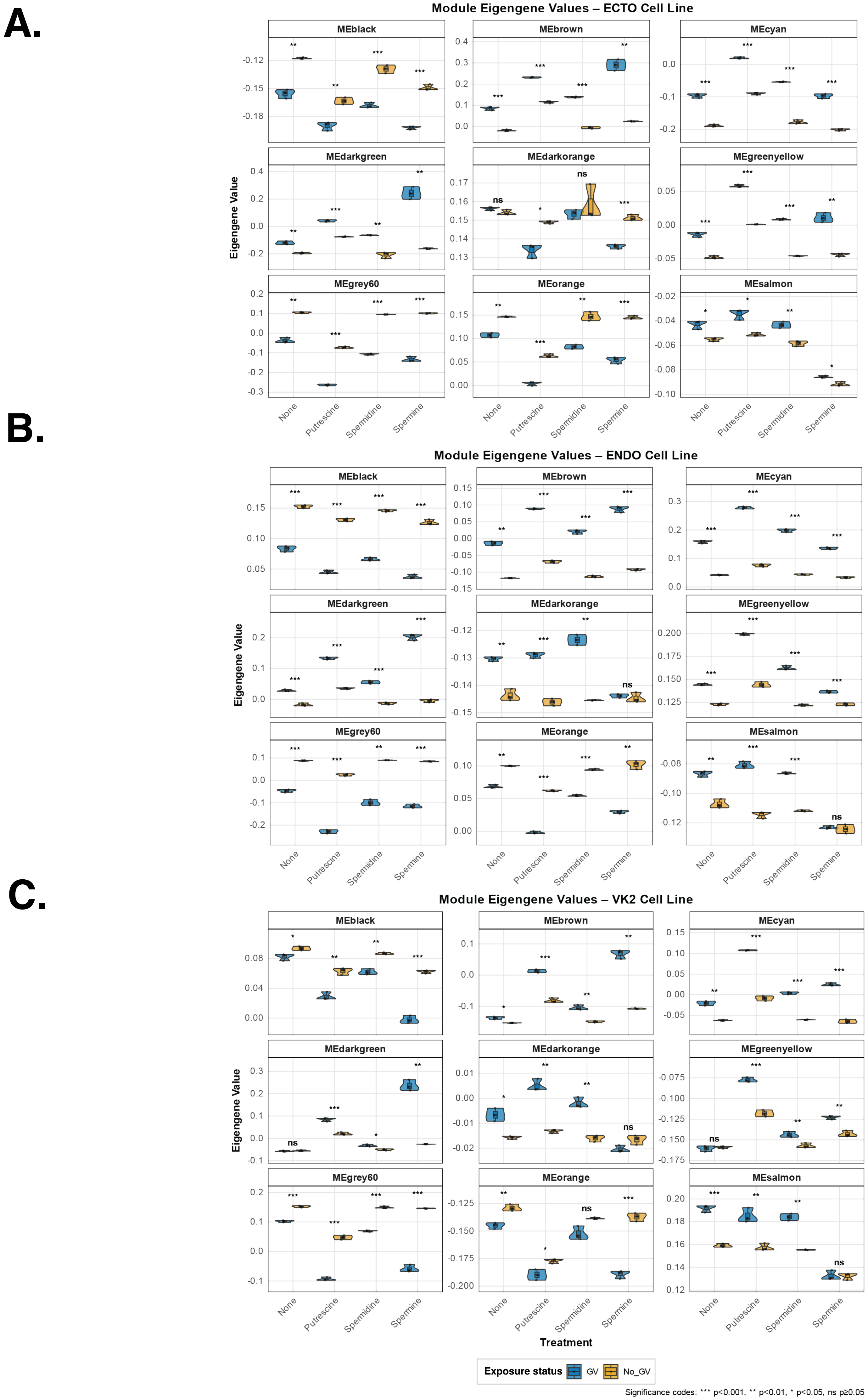
